# Incipient signs of genetic differentiation among African elephant populations in fragmenting miombo ecosystems in south-western Tanzania

**DOI:** 10.1101/210179

**Authors:** Alex L. Lobora, Cuthbert L. Nahonyo, Linus K. Munishi, Tim Caro, Charles Foley, Colin M. Beale, Lori S. Eggert

**Author notes:** Corresponding author: Alex L. Lobora, Address: Tanzania Wildlife Research Institute, P.O Box 661, Arusha, Tanzania.

## Abstract

Habitat fragmentation plays a major role in the reduction of genetic diversity among wildlife populations. The African savannah elephant population of the Ruaha-Rungwa and Katavi-Rukwa ecosystems in south-western Tanzania, comprises one of the world’s largest remaining elephant populations, but is increasingly threatened by loss of connectivity and poaching for ivory. We investigate whether there are incipient signs of genetic isolation (loss of heterozygosity) within the younger cohort as a result of habitat loss between the two ecosystems. To investigate the genetic structure of populations, we compared the genotypes for 11 microsatellite loci in the western (n = 81 individuals from Katavi-Rukwa), central (n = 36 individuals from Lukwati and Piti), and eastern populations (n = 193, individuals from Ruaha-Rungwa). We found evidence of significant genetic differentiation among the three populations, but the levels were low, suggesting recent divergence. Furthermore, we identified weak isolation by distance, suggesting higher gene flow among nearer individuals with samples within 50km of each other being more genetically similar to one another than beyond. Although sample sizes were small, a further analysis of genetic differences across populations and in separate age classes revealed evidence of increasing genetic structure among younger age classes across the landscape. In a long-lived species with overlapping generations, it takes a long time to develop genetic substructure even when there are substantial obstacles to migration. Thus, in these recently fragmented populations, inbreeding (and the loss of heterozygosity) may be less of an immediate concern than demography (the loss of adults due to illegal hunting).

## 1. Introduction

Human population growth is one of the main drivers of natural habitat loss and increased isolation of natural landscapes (DeClerck *et al*., 2010; Rands *et al*., 2010; Pereira *et al*., 2010). Habitat loss and fragmentation is a conservation problem not only because of the direct loss of range and increased edge effects (MacArthur & Wilson, 1967; Templeton *et al*., 1990; Murcia, 1995;Fischer & Lindenmayer, 2007; Hanski, 2011), but also because of the potential for inbreeding depression through genetic drift particularly in small isolated populations (Charlesworth *et al*., 1987; Frankham, 1996; Crnokrak *et al*., 1999; Hedrick & Kalinowski, 2000) which can increase extinction risk (Gibson *et al*., 2013). Moreover, the speed and scale at which fragmentation is happening is yet another cause of concern (Hansen *et al*., 2013; Haddad *et al*., 2015), because few migration routes are entirely within protected areas (Bolger *et al*., 2008; Harris *et al*., 2009; Bartlam-Brooks *et al*., 2011). A recent study conducted across five continents indicates that fragmentation of natural habitat reduces biodiversity by 13% to 75% with effects being greatest in the smallest and most isolated fragments (Haddad *et al*., 2015). Further reviews reveal that over the past half century, habitat destruction has led to the loss of more than a third of all forest cover worldwide (Hansen *et al*., 2013).

African elephant (*Loxodonta africana*) populations were historically distributed across many African countries (Douglas-Hamilton, 1987; Said *et al*., 1995; Barnes *et al*., 1999), with very little or no structure among populations because most populations were not isolated long enough (Georgiadis *et al*., 1994). But across their range fragmentation has been escalating, in particular within the past couple of decades, and populations are becoming increasingly restricted to protected areas (henceforth “PAs”) as a result of increasing human activity (Douglas-Hamilton *et al*., 2005; Graham *et al*., 2009; Jenkins & Joppa, 2009; Cantú-Salazar & Gaston, 2010; Craigie *et al*., 2010; UNEP, 2013). This poses a question of whether populations that were once widely ranging across the continent are becoming genetically isolated because of the ongoing habitat destruction and fragmentation. While it is important to recognize that there is a time lag between changes to habitats and the time when the full implications of those changes are experienced by wildlife species (Bennett,1999), it is desirable to understand early signs of variation among populations through the detection of genetic differentiation. Conservation genetics offers an ideal tool for this kind of investigation given the power it possesses in detecting differentiation of populations (Pritchard *et al*., 2000; Sunnucks, 2000, 2001; Rousset 2001a, b; Frankham *et al*., 2002; Manel *et al*., 2003; Broquet *et al*., 2006; Taylor *et al*., 2011; Paule *et al*., 2012). Information contained in a series of individual genotypes can quantify the extent to which isolated populations have lost genetic diversity over time, making it a relevant tool for assessing differences in structure within and among populations of the same species in fragmenting habitats (Taylor *et al*., 2011).

The past 20 years has seen widespread deforestation of the miombo woodlands in areas between Katavi-Rukwa and Ruaha-Rungwa ecosystems in southwestern Tanzania, with about 17.5% of the woodlands and forests modified or removed to make way for agricultural development, threatening connectivity between these ecosystems (Lobora *et al.*, 2017). This has clear implications for the viability of wide ranging species such as African elephants, for which one of the world’s largest remaining populations (about 12,000 individuals) is found in the two ecosystems combined (TAWIRI, 2014; Chase *et al*., 2016). Because fragmentation in this landscape is relatively recent (Lobora *et al*., 2017), and because elephants are long-lived (Armbruster, 1993) and are known to show little structure across the continent (Comstock *et al*., 2002), we expect there to be no structure among adult populations at evolutionary time scales across the landscape. However, if adult movement has recently become restricted due to the recent fragmentation of the Katavi-Rukwa and Ruaha-Rungwa ecosystems, we expect to see incipient signs of genetic structure, particularly among the younger cohorts within these populations.

## 2. Material and methods

### 2.1 Study area

The study area covers about 109,050 km^2^ and lies between latitude 6°15'59.38" to 8°10'23.78" S and longitude 30°45'13.29" to 35°28'34.44" E and comprises the Katavi-Rukwa ecosystem in the west, a contingent of Game Reserves (henceforth “GRs”), Game Controlled Areas (GCAs) and Open Areas (OAs) in the central part, as well as the Ruaha-Rungwa ecosystem in the east (Figure 1). About 45,961 km^2^ of this area is designated as Fully Protected Areas (Two National Parks-NPs, Seven GRs where minimal human activities are permitted), and 34,196 km^2^ designated as Lesser Protected Areas (Eight GCAs and Eight OAs where human activities are permitted alongside wildlife conservation). A further 28,893 km^2^ of land within the study area is unprotected and includes towns and highly populated regions north and south of Katavi National Park, and to the north-east and south of Ruaha National Park (Figure 1).

**Figure 1:**
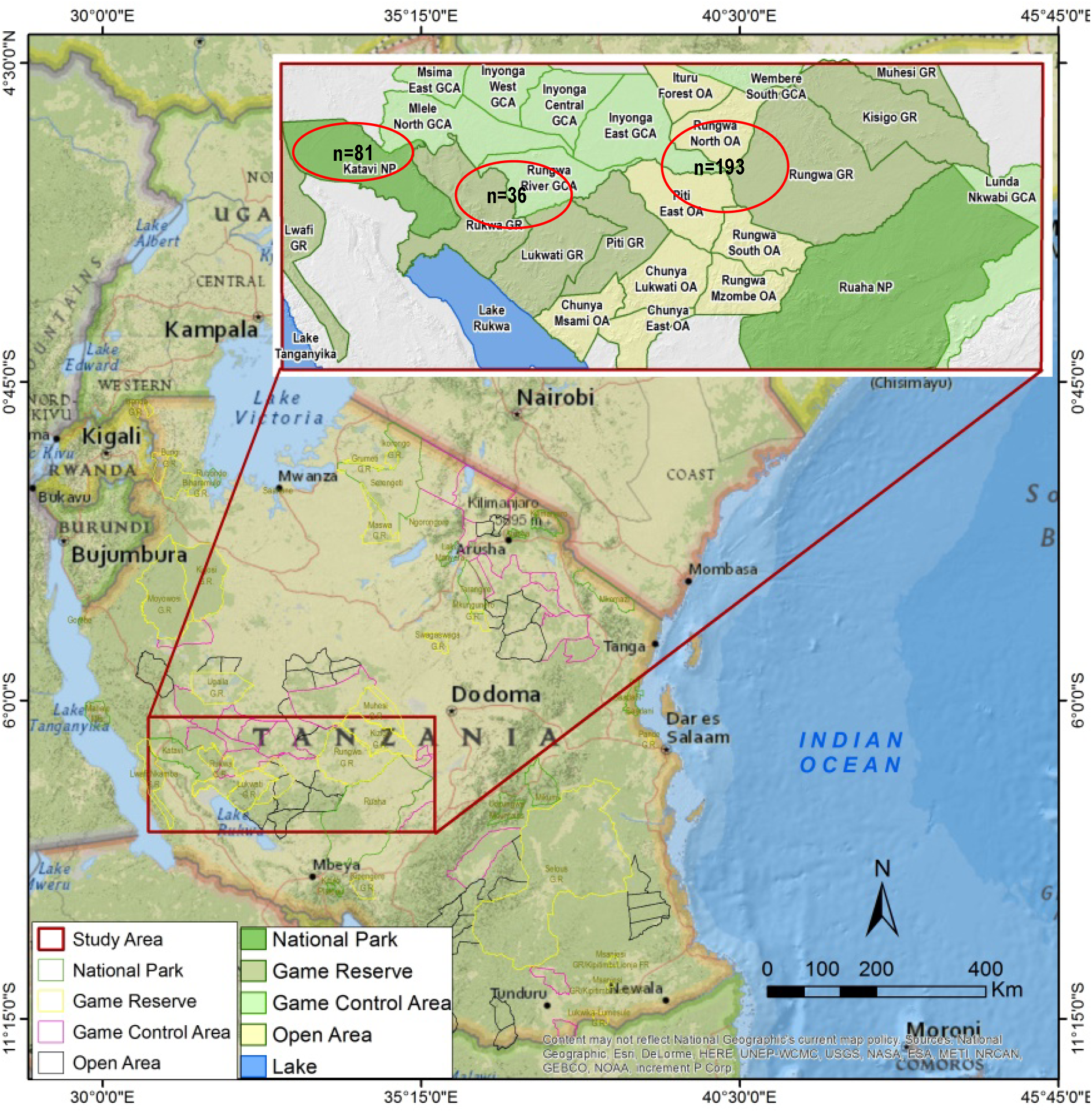
Sampling locations across the study area

### 2.2 Methods

#### Sample collection

We collected 380 fresh dung samples between July and November 2015 in Katavi-Rukwa ecosystem (henceforth “western population”), Lukwati and Piti Game Reserves (henceforth “central population”) and Ruaha-Rungwa ecosystem (henceforth “eastern population”). We employed an opportunistic random sampling strategy to obtain samples from different parts of the study area whilst avoiding samples from closely related individuals (e.g. in the event that a group of fresh samples were encountered in the same location, we only collected one sample). For each sample, we placed approximately 10 g of the external region of the dung bolus surface with genetic content (< 12 hours old) in 40 ml polypropylene tubes and boiled them for 15 min in the field to stall microbial activity, then preserved in Queens College Buffer (20% DMSO, 100 mM Tris pH 7.5, 0.25 M EDTA, saturated with NaCl; Amos *et al*., 1992). Samples were initially kept samples in the dark at room temperature in the field station and later moved them to a lab at the Nelson Mandela African Institution for Science and Technology (NM-AIST) for post field storage and subsequently shipped to the University of Missouri-Division of Biological Sciences under USDA permit number 128686 for subsequent DNA extraction and analyses.

#### DNA extraction, PCR, Sexing and microsatellite genotyping

We used the QIAamp mini stool extraction kit (Qiagen, Valencia, CA) to extract DNA from samples following earlier published protocols (Archie *et al*., 2006). The extraction process took place in a laboratory designated exclusively for the extraction of DNA from non-invasively collected samples to minimize the possibility of contamination (Okello *et al*., 2008; Ahlering *et al*., 2011). We genotyped all samples at 11 dinucleotide microsatellite loci developed for the African elephant (FH1, LaT24, FH60, LA5, FH19, LafMS06, LA6, LaT08, LafMSO2, FH48 and FH67), using published primers (Nyakaana *et al*., 2001; Archie *et al*., 2006; 2008, Eggert *et al*., 2008; Okello *et al*., 2008; Kongrit *et al*., 2008) with fluorescent labels. Multiplex PCR reactions (Ahlering *et al*., 2011) were performed using Platinum Multiplex PCR Master Mix (Applied Biosystems, Foster City, CA) following the manufacturer protocols, but in 8 µl volumes with 0.8X BSA and GC enhancer solution added to a final concentration of 10%. The PCR profile included an initial denaturing step at 95 °C for 2 min, followed by 40 cycles of 95 °C denaturing for 30 s, annealing at locus specific temperatures for 90 s, and 72 °C extension for 60 s; and a 30-min extension at 60 °C. A negative control was included in each PCR plate to detect contamination of the PCR reagents and a positive control sample was included to standardize scoring. We genotyped all samples on an ABI 3730XL capillary sequencer and subsequently analyzed with GeneMarker v2.6.7 (Soft Genetics LLC). To minimize the probability of genotyping error, we repeated the matching of both heterozygotes and homozygotes three times (Frantz *et al*., 2003; Hansen *et al*., 2008; Ahlering *et al*., 2011).

We used the Excel Microsatellite Toolkit (Park, 2001) to identify potential genotyping errors, create input files for population genetic analysis programs, find duplicates or genetically identical samples and calculate allele frequencies and diversity statistics. Because DNA extracts from non-invasively collected samples are dilute and contain degraded DNA, we rechecked each pair of genotypes that differed at 3 or fewer loci for possible problems with allelic dropout and considered genotype to represent the same individual if they differed at two or fewer alleles but matched in sex and had very similar bolus circumferences (Ahlering *et al*., 2011). This conservative approach was taken to avoid scoring samples as different individuals when they are actually erroneous genotypes (Ahlering *et al*., 2011).

To determine individual sex, we followed Ahlering *et al*. (2011). PCR concentration constituted 25 µl reactions mixture containing 0.1 U AmpliTaq Gold DNA Polymerase (Applied Biosystems), 1.25µl PCR Gold Buffer (Applied Biosystems), 1.25 µM dNTPs, 0.50 µl SRY1 forward primer, 0.5 µl SRY1 reverse primer, 0.5 primer, 0.5 µl AMELY2 reverse primer, 0.5 µl PLP1 forward primer, 0.5 µl PLP1 reverse primer, 1.00 µl MgCl2, 1.00 µl BSA and 1 µl DNA extract, 7 µl of the amplification products. PCR consisted of an initial denaturation at 95°C for 10 min, followed with 45 cycles of 30 sec at 95°C for 30 sec, at 59°C for 30 sec and 45 sec at 72°C, with a final extension of 10 min at 72°C with each PCR plate containing a negative and positive controls to detect for possible contamination of the PCR product and consistency of the amplification respectively (Ahlering *et al*., 2011). About 5 µl of PCR product was subsequently electrophoresed at 80 V for 40 min on a 2% Agarose gel stained with Gel Star. Since the restriction site is on the Y-chromosome, we scored single bands as females and three bands as males and repeated the process once for each sample to confirm sex (Ahlering *et al*., 2011).

## 3. Genetic analysis

We analyzed the set of unique genotypes within and among populations using GenePop 4.2 (Raymond & Rousset 1995, Rousset, 2008) to test for deviations from expected heterozygosity values under Hardy-Weinberg equilibrium (HWE), for linkage disequilibrium, and to determine the number of alleles at each locus (A), the observed (H_O_) and expected (H_E_) heterozygosity values, and the coefficient of inbreeding (F_is_) as estimated by Weir & Cockerham, 1984. Because sample sizes were unequal, we also used rarefaction in HP-Rare (Kalinowski, 2005) to estimate allelic richness, i.e. the mean number of alleles at a sample size of 36 (the smallest sample size for any population). We compared rarefied allelic richness among populations using ANOVA. We estimated genetic distances (fixation index- F_st_) between pairs of populations in Arlequin version 3.5.1.3 (Excoffier & Lischer 2010), and evaluated the significance of these F_st_ using a permutation test (1000 permutations). Finally, we estimated the number of migrants (*Nm*) among pairs of populations indirectly based on genetic data through F_st_ using the equation:

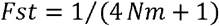

Where *N* represents effective population size of each population and *m* is the migration rate between populations.

We tested for genetic differentiation using F_st_ across all individuals in the three populations and then compared F_st_ across groups of different age cohorts i.e. Juvenile age 0-3, Young age 4-9, Sub-adult age 10-19 and Adult age 20+. We obtained the age structure of the three populations through dung bolus measurements following Morrison *et al.* (2005). We then summarized the large number of possible comparisons of fixation index across different age cohorts over the three populations and subsequently counted the number of significant differences relative to all possible combinations of those age-classes (landscape comparisons). For example, adult elephants in the eastern population were compared to the three sub-adult elephant populations (within the eastern, and against both other populations) resulting in any inter-age class comparison having a total of nine possible significant comparisons, and intra-age class comparisons having a total of three comparisons.

To investigate possible patterns of isolation-by-distance (IBD), we used an individual-based approach. We computed a pairwise matrix of inter-individual genetic distances using the Bray-Curtis percentage dissimilarity measure (function *diss.dist* from the R-package *poppr;* Kamvar *et al.,* 2014) that we compared to the corresponding pairwise matrix of inter-individual Euclidean distances using a simple Mantel test with 10000 permutations (function mantel.randtest from the R-package ade4: Dray & Dufour, 2007). Additionally, based on the geographic coordinates of sample locations, we investigated spatial patterns of IBD using a Mantel autocorrelogram in the MATLAB software-coding environment (Mathworks, Inc.; Prunier *et al.,* 2013). Euclidean distance classes were defined every 50,000 m (up to 50 km), resulting in 10 binary matrices representing the membership of genetic distances to the distance class being tested (with 0 for pairs of individuals belonging to the same distance class and 1 otherwise). Each binary matrix was compared to the genetic distance matrix using a simple Mantel test with 1,000 permutations. We then plotted Mantel correlation values over distance classes, with a 95 % confidence interval determined by bootstrap resampling (1,000 iterations and random removal of 20% of individuals at each iteration). Mantel autocorrelograms were also computed for each sex separately.

To test for structure on an evolutionary timescale, we analyzed genotypes in STRUCTURE 2.3.4 (Pritchard *et al*., 2000), denoting a model-based clustering algorithm to multi-locus genotype data (Ishida *et al*., 2016). We programmed length of the burn-in period to 10,000 and the number of Markov Chain Monte Carlo (MCMC) reps after the burn-in to at least 100,000 steps. We further programmed STRUCTURE to run 10 times for each value of K from 1 to 10, with the use of prior information about the location from which the sample was collected, under the admixture-correlated model and correlated allele frequencies among populations. We further used spatial Principal Component Analysis (Jombart et *al.,* 2008) to reveal possible cryptic genetic structures, stemming from the specific life-history traits of this long-lived species. To that aim, we used a distance-based neighborhood network with a distance threshold consistent with results from the Mantel correlograms. Global and local Monte Carlo tests were carried out with 10,000 permutations to evaluate the significance of detected patterns (Jombart *et al.,* 2008).

## 4. Results

Of the 380 samples collected across the landscape, 376 (98.9%) were successfully genotyped and 310 individuals identified by their unique genotypes. The remaining 66 samples were recaptures within the same populations and therefore discarded from subsequent analyses. The age class distribution by sex of the three populations is presented in Table 1. We were unable to determine sexes for some individuals due to repeated failure to discriminate bands as either males or females.

**Table 1:** Age class distribution by sex

| **Sex** | **Adult** | **Sub adult** | **Young** | **Juvenile** | **Unknown** |
| --- | --- | --- | --- | --- | --- |
| **Female** | 68 | 41 | 11 | 0 |  |
| **Males** | 90 | 39 | 8 | 1 |  |
| **Unknown** |  |  |  |  | 52 |
|  | **158** | **80** | **19** | **1** | **52** |

Two of the 11 loci (LafMS02 and LafMS06) did not conform to expectations under HWE in any of the three populations after applying Bonferroni correction for multiple tests (Rice, 1989). These loci had significant excesses of heterozygosity that could not be resolved through reanalysis of the genotypes and hence were removed from the analyses. Other than these loci, LA5 and FH19 deviated from expectations in the western population, FH60 deviated in the central population, and LA5 and FH48 deviated in the eastern population. Because there were no consistent patterns of deviation across populations, these loci were retained in the analyses (Table 2).

**Table 2:** Genetic diversity measures for the populations sampled for this study. A=number of alleles detected, A_R_=number of alleles adjusted for unequal sample sizes, A_P_=number of private alleles, H_O_=observed heterozygosity, H_E_=expected heterozygosity, F_IS_=coefficient of inbreeding, not conform to expected values under Hardy-Weinberg equilibrium

| Population | Locus | N | A | $A_R$ | $A_P$ | $H_O$ | $H_E$ | $F_{IS}$ |
| --- | --- | --- | --- | --- | --- | --- | --- | --- |
| **Western**<br>**n=81** | FH1 | 80 | 5 | 4.2 | 0.1 | 0.675 | 0.687 | 0.018 |
|  | LaT24 | 80 | 10 | 7.7 | 1.0 | 0.688 | 0.829 | 0.171 |
|  | FH60 | 81 | 5 | 3.0 | 0.1 | 0.296 | 0.330 | 0.102 |
|  | LA5 | 70 | 7 | 5.8 | 1.5 | 0.429* | 0.667 | 0.359 |
|  | FH19 | 80 | 9 | 6.6 | 1.2 | 0.513* | 0.712 | 0.282 |
|  | LA6 | 81 | 7 | 4.2 | 0.9 | 0.605 | 0.585 | -0.035 |
|  | LaT08 | 79 | 10 | 7.7 | 0.4 | 0.696 | 0.842 | 0.174 |
|  | FH48 | 81 | 9 | 6.8 | 0.3 | 0.802 | 0.726 | -0.106 |
|  | FH67 | 81 | 8 | 4.6 | 0.2 | 0.469 | 0.598 | 0.197 |
|  | Average |  | 7.8 | 5.6 | 0.6 | 0.605 | 0.664 | 0.129 |
|  | StDev |  | 1.9 | 1.7 | 0.5 | 0.170 | 0.153 | 0.150 |
| **Central**<br>**n=36** | FH1 | 36 | 4 | 4.0 | 0.0 | 0.750 | 0.720 | -0.042 |
|  | LaT24 | 36 | 7 | 6.5 | 0.1 | 0.694 | 0.812 | 0.146 |
|  | FH60 | 36 | 4 | 3.4 | 0.3 | 0.306* | 0.504 | 0.397 |
|  | LA5 | 29 | 4 | 3.6 | 0.0 | 0.345 | 0.482 | 0.288 |
|  | FH19 | 36 | 7 | 5.9 | 0.5 | 0.528 | 0.735 | 0.288 |
|  | LA6 | 36 | 4 | 3.7 | 0.6 | 0.528 | 0.570 | 0.075 |
|  | LaT08 | 36 | 10 | 9.1 | 1.2 | 0.750 | 0.826 | 0.094 |
|  | FH48 | 36 | 8 | 7.2 | 0.3 | 0.750 | 0.724 | -0.037 |
|  | FH67 | 36 | 8 | 6.7 | 1.0 | 0.694 | 0.730 | 0.049 |
|  | Average |  | 6.2 | 5.6 | 0.4 | 0.630 | 0.678 | 0.140 |
|  | StDev |  | 2.3 | 2.0 | 0.4 | 0.148 | 0.128 | 0.154 |
| **Eastern**<br>**n=193** | FH1 | 187 | 7 | 4.6 | 0.6 | 0.706 | 0.678 | -0.041 |
|  | LaT24 | 190 | 11 | 7.6 | 1.0 | 0.763 | 0.826 | 0.076 |
|  | FH60 | 193 | 5 | 4.0 | 0.9 | 0.368 | 0.422 | 0.128 |
|  | LA5 | 170 | 8 | 5.2 | 0.9 | 0.535* | 0.664 | 0.194 |
|  | FH19 | 187 | 10 | 6.8 | 1.2 | 0.642 | 0.737 | 0.130 |
|  | LA6 | 192 | 8 | 3.7 | 0.5 | 0.625 | 0.556 | -0.124 |
|  | LaT08 | 184 | 11 | 7.5 | 0.4 | 0.788 | 0.827 | 0.047 |
|  | FH48 | 192 | 9 | 6.8 | 0.3 | 0.922* | 0.759 | -0.215 |
|  | FH67 | 193 | 9 | 6.4 | 0.7 | 0.674 | 0.705 | 0.044 |
|  | Average |  | 8.7 | 5.9 | 0.7 | 0.652 | 0.686 | 0.027 |
|  | StDev |  | 1.9 | 1.5 | 0.3 | 0.139 | 0.130 | 0.131 |

### 4.1 Genetic diversity within populations

We found high levels of genetic diversity in all populations (Table 2), with allelic diversity ranging from an average of 8.7 (± 1.9 StdDev) alleles per locus in the eastern population to 6.2 (± 2.3 StdDev) alleles per locus in the central population. When these values were corrected to a standard sample size of 36 (the size of the smallest sample) using rarefaction (Kalinwoski, 2004), there was no significant difference among populations in the number of alleles (ANOVA, df=2, p=0.703) or private alleles (ANOVA, df=2, p=0.408).

### 4.2 Genetic differentiation among populations

We found the three populations to be significantly different, though the level of differentiation was small (Table 3), with F_st_ values ranging from 0.006 between the eastern and central populations to 0.011 between the western and central populations. The estimated number of migrants ranged from 22.5 individuals per generation between the western and central populations to 41.4 individuals per generation between the eastern and central populations.

**Table 3:**
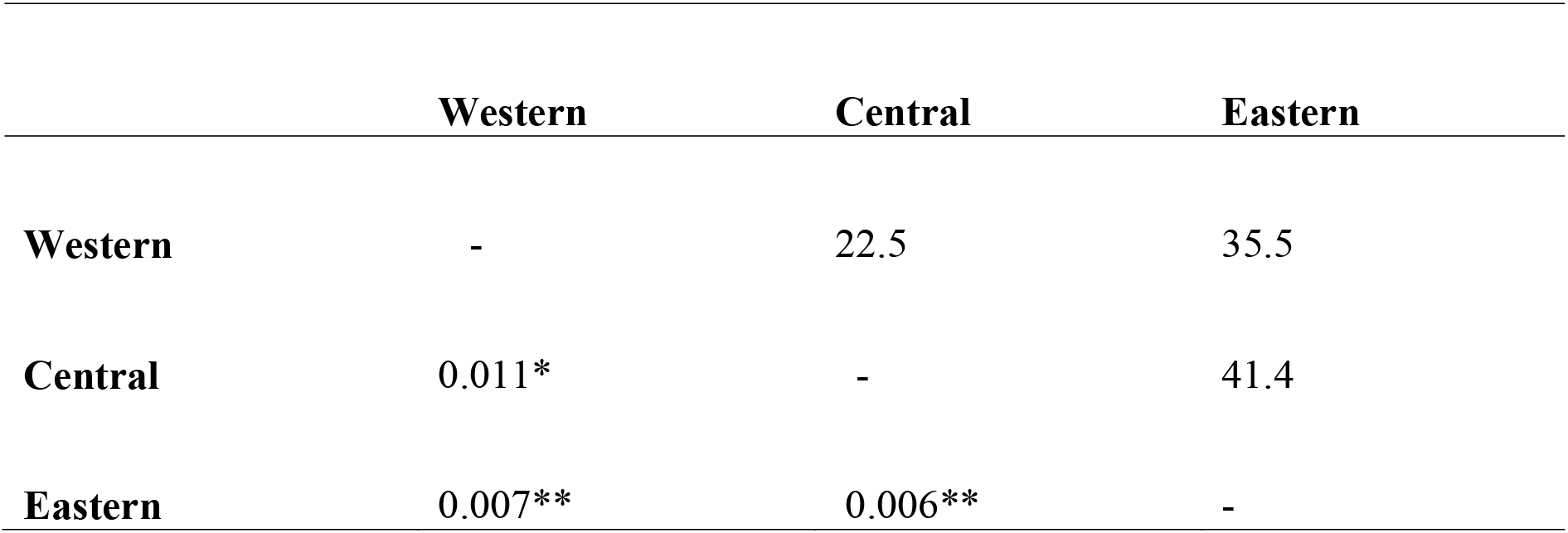
Genetic distance measures among populations are shown below the diagonal. Significance levels are indicated as *=p<0.05, **=p<0.01. The number of estimated migrants per generation is shown above the diagonal.

|  | Western | Central | Eastern |
| --- | --- | --- | --- |
| Western | - | 22.5 | 35.5 |
| Central | 0.011* | - | 41.4 |
| Eastern | 0.007** | 0.006** | - |

We identified weak (but statistically significant) isolation by distance (Mantel test, r = 0.09, p = 0.045), suggesting higher gene flow among nearer individuals. Samples within 50km of each other were more likely to be genetically similar to one another, but beyond this distance there was no remaining population structure (Figure 2a). This IBD pattern hold when considering females only (Figure 2b), whereas, genetic relatedness among males was not significant for the first 100 km, suggesting high dispersal propensity in this sex (Figure 2c).

**Figure 2:**
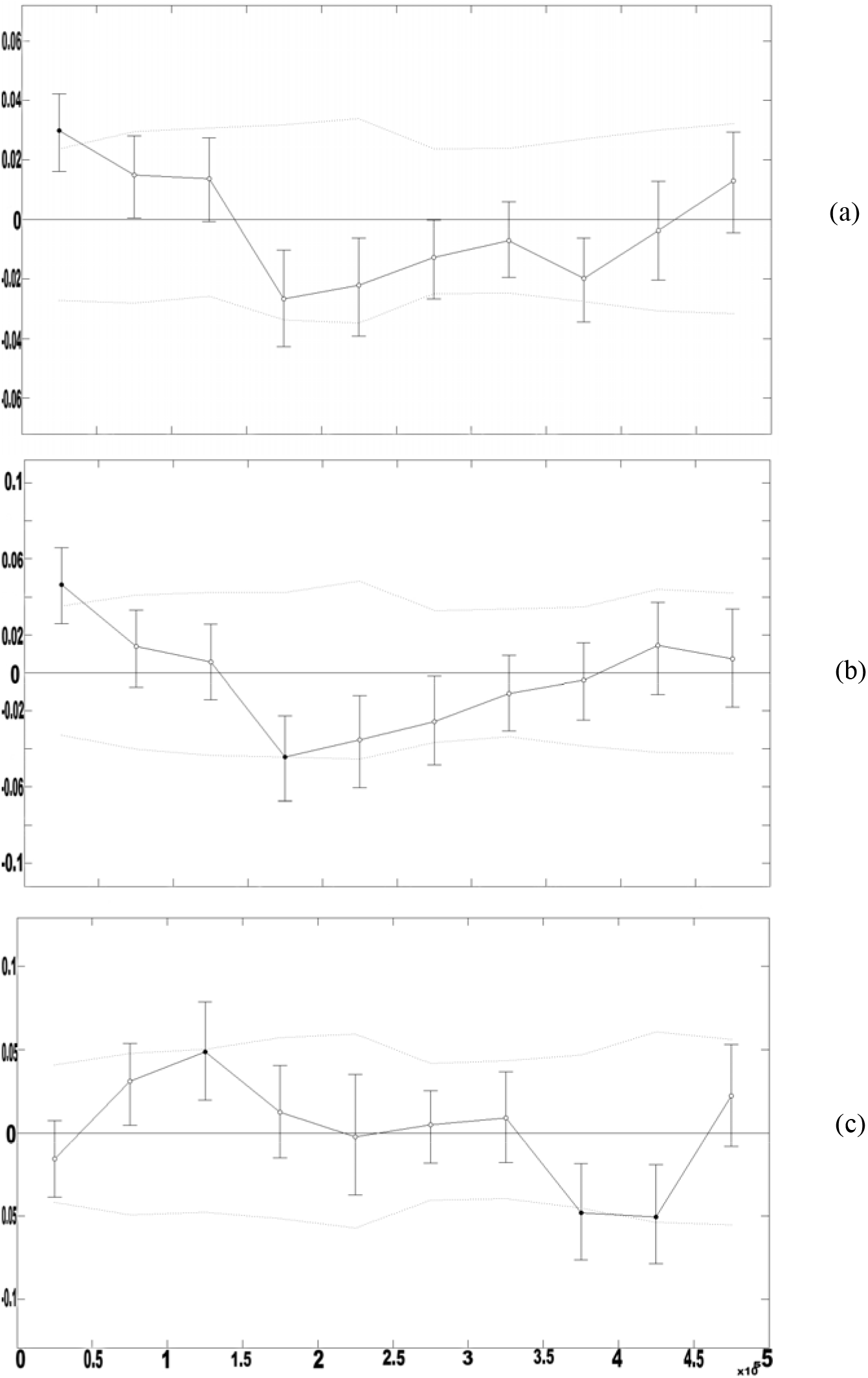
Spatial patterns of genetic autocorrelation (a) among all individuals, (b) among females and (c) among males. r: standard Mantel correlation with 1000 permutations. Error bars bound the 95% confidence interval about r as determined by bootstrap resampling. Upper and lower confidence limits (dotted line) bound the 95% confidence interval about r under the null hypothesis of no spatial structure. Black dot indicates significant Mantel test at 5%.

Analyses in STRUCTURE detected no significant genetic clustering among populations across the study landscape (K=1), suggesting that while there is significant differentiation over a recent, ecological timescale (Table 2), individuals represent a single genetic population over an evolutionary timescale. Nevertheless, we identified significant cryptic genetic structures when using sPCA. The global Monte-Carlo test performed in sPCA was significant (max(t) = 0.007, p = 0.007), indicating the presence of a significant global genetic structure. On the contrary, the local Monte-Carlo test did not detect any significant local structure (max(t) = 0.013, P > 0.05). Scores of individuals along the first sPCA axis distinguished the western population from the central and eastern populations (Figure 3a), in accordance with the numbers of migrants previously estimated (Table 3). Along the second axis though, individuals showed a spatial pattern characterized by a longitudinal alternation of genetic clusters, roughly delimited every 50 km, highlighting the influence of a continuous IBD in this species (Figure 3b).

**Figure 3:**
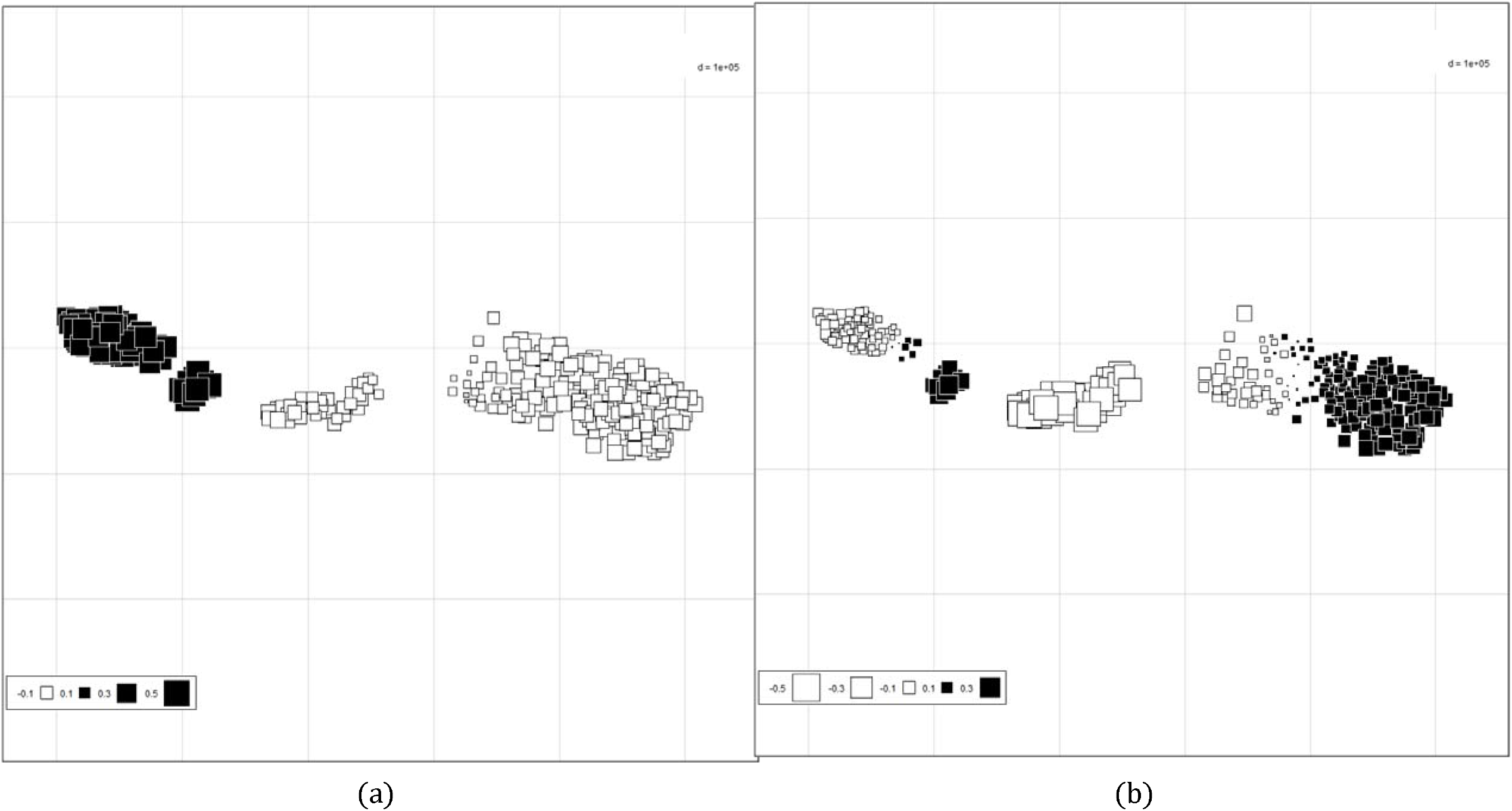
Maps of the first (a) and second (b) global sPCA scores. Large white and black squares stand for highly negative and positive scores respectively. Small squares stand for low sPCA scores.

### 4.3 Genetic differentiation among age cohorts

Results of comparisons of fixation index across different age classes and over the three populations shows just one (of three possible) significant F_st_ difference for each intra-age comparison (Table 4), but found strong evidence that F_st_ differences increase across age classes: We found two (of possible nine) adult to sub-adult comparisons to be significant, and five (of possible nine) adult to younger cohort comparisons to be significant, suggesting growing differences among the younger cohort in the population across the landscape (Table 4).

**Table 4:** Genetic distance measures among age groups. Significance levels are indicated as *=p<0.05, **=p<0.01, ***p<0.001. Negative values denote genetic distances not significantly different from zero whereas A, S, Y denotes Adults, Sub adults and Young populations respectively.

|  | Western A | Western S | Western Y | Central A | Central S | Central Y | Eastern A | Eastern S | Eastern Y |
| --- | --- | --- | --- | --- | --- | --- | --- | --- | --- |
| Western A | - |  |  |  |  |  |  |  |  |
| Western S | 0.002 | - |  |  |  |  |  |  |  |
| Western Y | -0.002 | -0.013 | - |  |  |  |  |  |  |
| Central A | 0.001 | 0.009 | -0.011 | - |  |  |  |  |  |
| Central S | 0.013 | 0.020* | 0.006 | -0.006 | - |  |  |  |  |
| Central Y | 0.034** | 0.035* | 0.029 | 0.035* | 0.003 | - |  |  |  |
| Eastern A | 0.009*** | 0.006** | 0.000 | 0.007 | 0.007 | 0.028* | - |  |  |
| Eastern S | 0.006* | 0.005 | 0.001 | -0.003 | 0.001 | 0.020* | 0.001 | - |  |
| Eastern Y | 0.018* | 0.013 | 0.004 | -0.008 | 0.014 | 0.043* | 0.015* | -0.001 | - |

## 5. Discussion

We found evidence of weak but significant genetic differentiation among the three recently divided populations, particularly between younger elephants, suggesting that the recent loss of natural habitat (Lobora *et al*., 2017), may be starting to generate population level differences (Table 3). Isolation was weakly but significantly associated with distance in the population at a whole and for females, consistent with early stages of population fragmentation. Landguth *et al*. (2010) found the lag time to barrier detection with genetic methods to be relatively short (1–15 generations) for wide ranging species but cautioned that detecting the effects of fragmentation on long-lived species (with overlapping generations) over ecological time scales may be difficult. Thus, we are not surprised that STRUCTURE did not detect significant genetic clustering. Although this program works well when population structure is relatively weak (Hubisz *et al*., 2009), it may fail to detect structure when differentiation levels are as low as those in this study (Duchesne & Turgeon, 2012). This is also consistent with our prediction that there would be no significant structure across adult individuals in these populations at evolutionary time scales because habitat fragmentation is a recent phenomenon. Nevertheless, sPCA revealed subtle global hierarchical genetic structure, with eastern and central populations showing higher genetic relatedness than the western population at the higher level of hierarchy, confirming inferred migration rates. This is unsurprising because habitat loss/fragmentation due to anthropogenic activities is higher between western and central populations than between central and eastern populations (Lobora *et al*., 2017). At the lower level of the hierarchy, it appeared that genetic structuring mostly stemmed from a longitudinal IBD pattern, with a lag distance of about 50 km, suggesting that IBD is an important driver of genetic differentiation in this system.

The historical large extent of miombo woodland linking these three populations appears to have facilitated broad-scale connectivity, at least until recently. Our recent analysis on the broad area extending from the Ruaha-Rungwa ecosystem to the Katavi-Rukwa ecosystem indicates that these areas retained approximately 73% of miombo woodland cover up until 1990s and continuous connectivity may only have been impaired recently (Lobora *et al*., 2017). Despite large areas of natural woodland remaining between the two ecosystems even now habitat loss has limited movement between the two ecosystems to a very narrow region (corridor), including some areas heavily used by people and a main road linking the northern and southern regions of Tanzania (Caro *et al*., 2009; Jones *et al.,* 2009).

The low level of genetic differentiation among populations could be partly explained by the fact that the average distance between farthest populations (about 200 km) is within the dispersal capabilities of African elephant (Blanc *et al*., 2007). The measure of population subdivision across all populations (F_st_), was low suggesting many successful migrants entering each population per generation (approximately 25 years for African elephant, Blanc, 2008) assuming an island model of migration (Frankham *et al*., 2002). On average across all populations, there were more than 20 successful migrants per generation (Table 3), a substantial level of gene flow among adults (Vucetich & Waite, 2000), suggesting that the cause of the small observed genetic differentiation is likely to be very recent. As confirmed by Mantel autocorrelograms (Figure 2b & 2c), dispersal of African elephants is sex-biased, and males typically disperse farther than females (Blake *et al.,* 2008; Petit & Excoffier, 2009), and gene flow by males can prevent serious losses of genetic diversity (Nyakaana & Arctander, 1999; Ishida *et al*., 2013). Without substantial levels of gene flow, habitat fragmentation and other anthropogenic disturbances can lead to extensive genetic differentiation among populations (Dixon *et al*., 2007), even among populations that are geographically close (Vos *et al*., 2001). However, our results were generated using an indirect method and should therefore be considered conservative and viewed with caution (Whitlock & McCauley, 1999).

We found no evidence of significant inbreeding in these populations, highlighting the importance of management actions (such as protection of the remaining potential habitat for connectivity) to maintain migration corridors that reinforce the gene flow. This is particularly important because conservation of wide-ranging species depends on PAs and dispersal areas to provide for protection and connectivity between them respectively (Western *et al*., 2009; Ahlering *et al*., 2012; Epps *et al*., 2013). Our analysis indicates fragmentation signs to be larger in the genetic structure of young individuals born when movement became increasingly restricted after 1990 and that genetic variations observed between adults and the young age could be precursors of what can be expected in the future. These changes will persist for at least a generation (even if a corridor were completely resurrected today) but appropriate management could restore a fully mixed population in the future.

Overall, the results obtained in our analysis are consistent with the suggestion that habitat fragmentation and loss constitute a threat to African elephant populations across their range (Comstock *et al*., 2002). As demonstrated in other taxa such as large carnivores (Johnson *et al*., 2001), African elephants are also susceptible to losses in genetic variation due to habitat fragmentation, despite long generation times (Blanc, 2008). The incipient signs of genetic differentiation detected in our analysis indicate increasing conservation challenges in human dominated landscapes (Newmark, 2008), calling for deliberate efforts and political will to save remaining dispersal areas for continued gene flow.

## Management Implications

A species’ ability to cope with the changing selective forces resulting from anthropogenic disturbance may be partially determined by the amount of genetic variability in populations as well as the way that variation is structured within and between populations (Archie *et al*., 2011; Ishida *et al*., 2016). Evidence for recent emergence of genetic structure within the three studied elephant populations suggests that habitat loss and fragmentation in the areas between Ruaha and Katavi are starting to alter population connectivity. At present a narrow corridor of natural habitat persists between the two systems, but heavy human use reduces the suitability of this for elephant movements. The remaining potential habitat for connectivity between the two ecosystems falls under the multiple landuse category (Open Areas), and we call for deliberate and timely actions to upgrade the protection status of this area to ensure continued gene flow between these populations. One of these efforts may include transforming these Open Areas (Piti east & Rungwa south) to a Wildlife Management Area (WMA), a new landuse category that promotes local community driven conservation allowing greater local community buy-in (Stoner *et al*., 2007; USAID, 2013; WWF, 2014). The current country political situation encourages WMA establishment over new National Parks and Game Reserves although in practice it can achieve the same goal if well implemented.

## Acknowledgements

We would like to thank the Ministries of Livestock development and Natural Resources and Tourism for facilitating the required permits for this study. We would also like to thank Mr. Amiri Swaleh for helping with sample collection.

## Supplementary materials

**Appendix 1:**
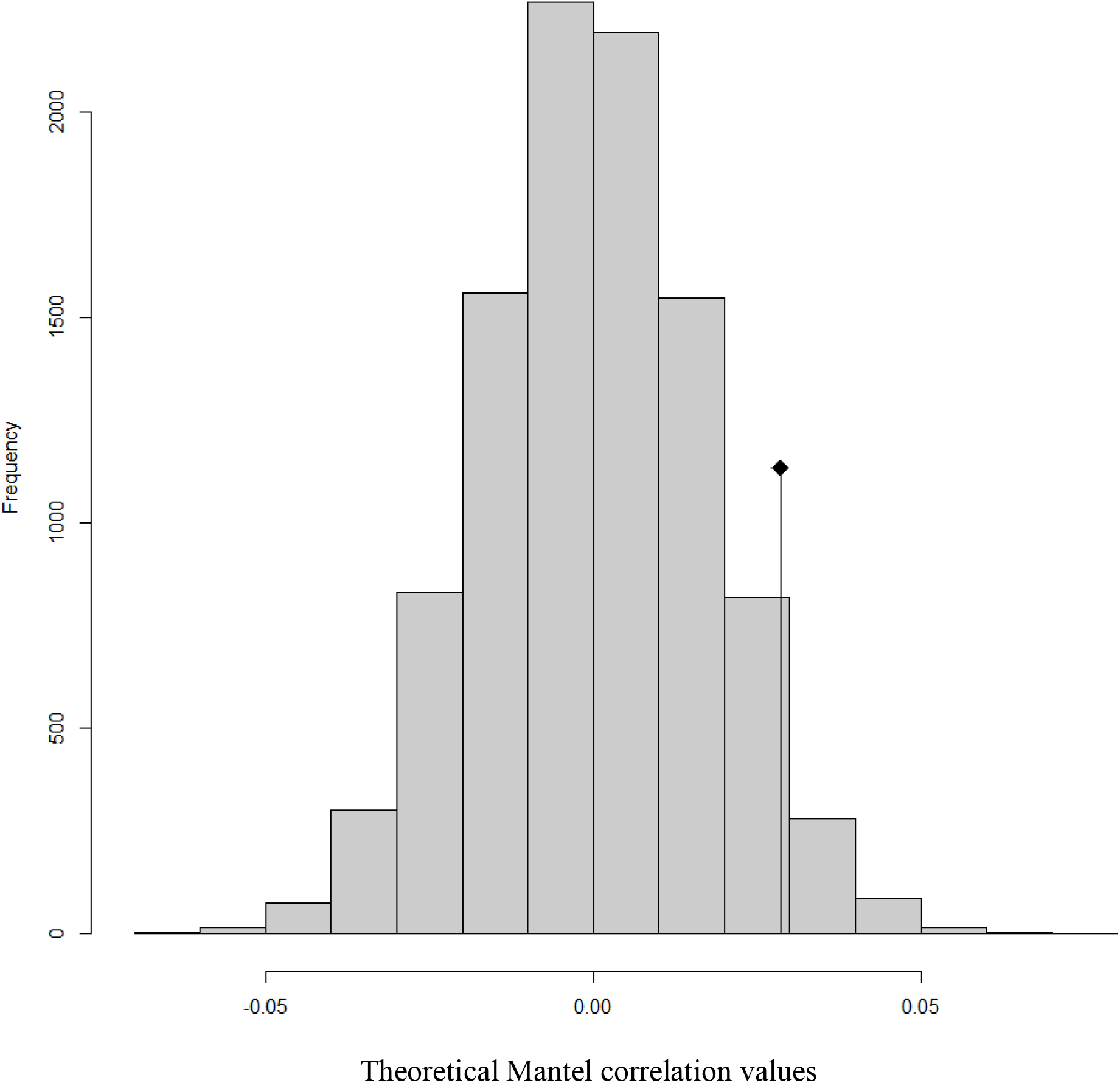
Theoretical distribution of Mantel correlation values under the null hypothesis. The vertical line with diamond indicates the observed Mantel correlation.

**Appendix 2:**
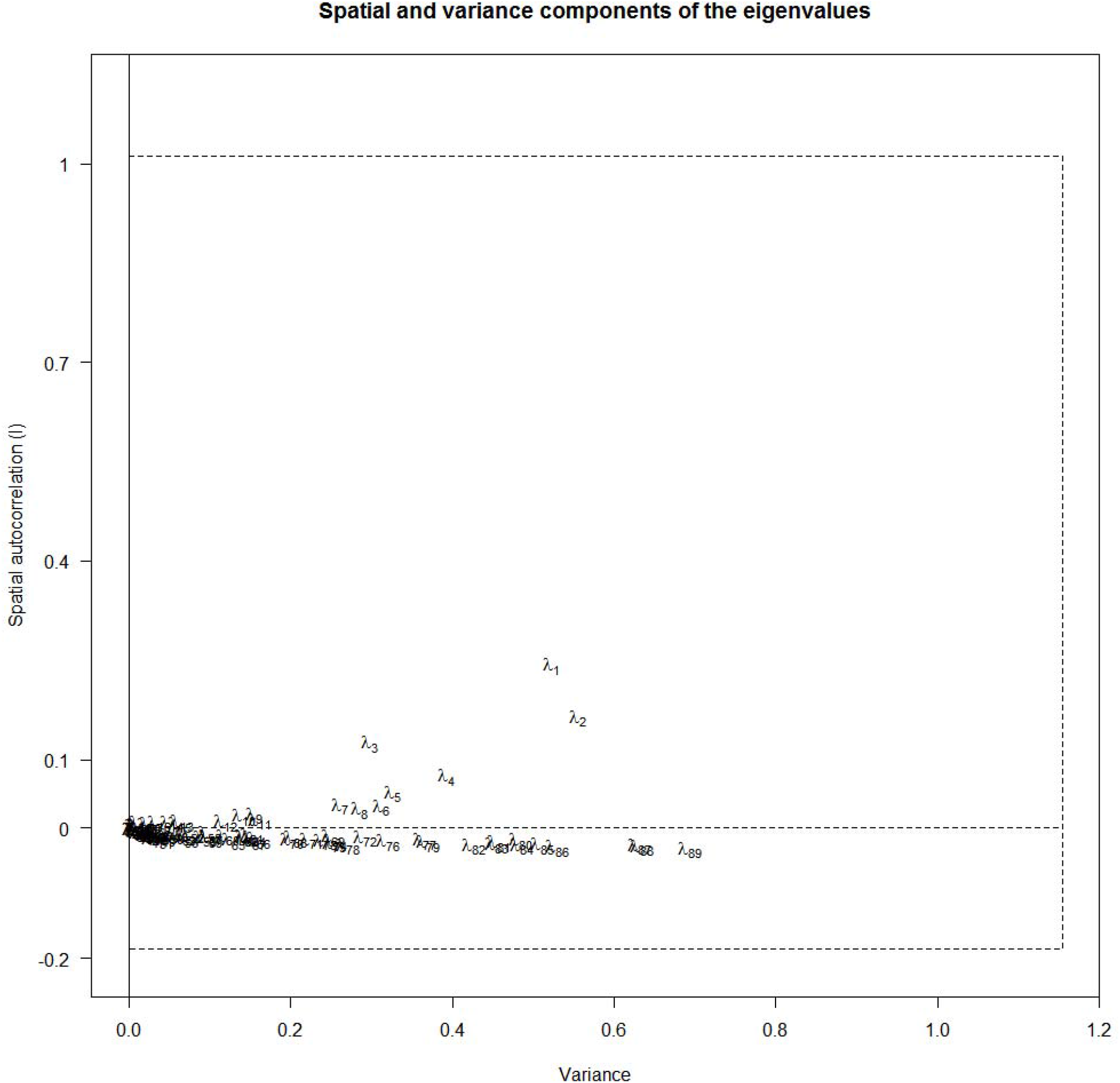
The respective importance of the two components of the sPCA (maximization of both variance and autocorrelation). Lambda values represent eigenvalues. It indicates that the two first principal components may be considered as meaningful as they stand for both a maximization of variance (0.6) and positive spatial autocorrelation (about 0.2). The dashed line stands for an absence of autocorrelation.

**Appendix 3**: Script used for the analysis

#Load libraries and functions

library(adegenet)

library(poppr)

library(ecodist)

Mat2Linear = function(x){

 x=data.matrix(x)

 dimm=dim(x)[1]

 if(sum(is.na(x[upper.tri(x)]))==((dim(x)[1]*(dim(x)[1]-1))/2)){

 x=t(x)}

 x=x[upper.tri(x, diag = FALSE)]

 x=as.data.frame(x)

} # this function is to convert data from matrix to dataframe format

#Prepare data in Genind format

setwd('C:/Users/User/Dropbox/UDSM/Data analysis/Chapter4/IBD')

R=read.structure("MP123.stru",n.ind = 310,n.loc = 11,onerowperind = TRUE,col.lab = 2,col.pop = 1,row.marknames = 1,col.others = 0)

tab(R, NA.method="mean")

Rxy=read.table("MP123_xy.txt",h=T)

Rtemp=data.frame(cbind(Rxy$x,Rxy$y))

R$other$xy=Rtemp

#Compute MATRICES and DATAFRAMES

DE=dist(x = R$other$xy,method = “euclidean”,diag = TRUE,upper = TRUE)

DG=diss.dist(R, percent = TRUE, mat = FALSE)

de=Mat2Linear(DE)

dg=Mat2Linear(DG)

dat=data.frame(dg=dg,de=de)

colnames(dat)=c('dg','de')

plot(dat$de,dat$dg)

abline(lm(dg~de,data=dat),col="red")

#dAPC on original populations

dapc1b=dapc(R,n.pca = 80,n.da = 2)

scatter(dapc1b,xax = 1,yax = 2,clabel = 0.5) # populations are not fully distinct

scatter(dapc1b,1,1, bg="white", scree.da=FALSE, legend=TRUE, solid=.4)

#dAPC with inference of the best number of populations

grp <- find.clusters(R,n.pca = 80,n.clust = 5) # choose n.pca=80 / choose n.clusters at minimum BIC : 5-7

table.value(table(pop(R), grp$grp), col.lab=paste("K", 1:11),row.lab=levels(R@pop))

dapc1a=dapc(R,pop = grp$grp,n.pca = 80,n.da = 3) # choose n.pca=80 / choose n.da (from 2 to n.da)

scatter(dapc1a,xax = 1,yax = 2,clabel = 0.5)

scatter(dapc1a,xax = 1,yax = 3,clabel = 0.5)

scatter(dapc1a,1,1, bg="white", scree.da=FALSE, legend=TRUE, solid=.4)

#IBD

ibd0 <- mantel.randtest(DG, DE,nrepet = 10000)

ibd0

plot(ibd0)

ibd=mantel(dg~de,data = dat,nperm = 10000)

ibd # pval3 provides pvalue for two-tailed test

mm=mgram(DG,DE,nclass = 10,nperm = 1000)

plot(mm,xaxt="n",ylim=c(-0.03,0.03))

axis(1, at=seq(0,max(dat$de),50000), labels=seq(0,max(dat$de),50000)) # significant positive autocorrelation at DE < 50000

#SPCA Analysis

spca1=spca(R,scannf = TRUE,type = 5,d1 = 0,d2 = 100000) # you can explore the effect of changing d2 (the scale of neighbourhood)

#HERE, you'll be asked to provide the number of positive components (e.g. 3), then the number of negative components (e.g. 0), according to provided graph

eigini=spca1$eig[c(1:5)]

eigscaled=eigini/eigini[1]

eigres=rbind(eigini,eigscaled)

eigres

loads <- spca1$c1[, 1]^2

names(loads) <- rownames(spca1$c1)

loadingplot(loads, xlab = “Alleles”, ylab = “Weight of the alleles”,main = “Contribution of alleles \n to the sPCA axis”)

s.value(Rxy, spca1$ls[, 1], origin=c(min(Rxy$x),min(Rxy$y)), csize = 0.5)

s.value(Rxy, spca1$ls[, 2], origin=c(min(Rxy$x),min(Rxy$y)), csize = 0.5)

s.value(Rxy, spca1$ls[, 3], origin=c(min(Rxy$x),min(Rxy$y)), csize = 0.5)

